# Revisiting Amplitude of Low-Frequency Fluctuations (ALFF) in Resting-state fMRI: Clarifications and Improvements

**DOI:** 10.1101/2025.08.07.669216

**Authors:** Lejian Huang, Rami Jabakhanji, Andrew D. Vigotsky, Paulo Branco, Marwan N. Baliki, A. Vania Apkarian

## Abstract

The amplitude of low-frequency fluctuations (ALFF) and its related measure, fractional ALFF (fALFF), are widely used resting-state fMRI techniques for quantifying spontaneous neural activity within specific frequency bands. However, inconsistencies in the definition and implementation of ALFF have led to confusion in the field. In this study, we provide a mathematical clarification of ALFF and fALFF by introducing two variants: the arithmetic mean-defined ALFF/fALFF (amALFF/amfALFF) and the quadratic mean-defined ALFF/fALFF (qmALFF/qmfALFF). We examine the relationships between mean BOLD intensity (MBI), amALFF, and qmALFF across both subjects and voxels using two independent datasets mapped onto different brain templates. Additionally, we investigate the impact of *z*-scoring the original BOLD signal on ALFF and fALFF metrics. Our key findings include: (1) MBI is positively correlated with both amALFF and qmALFF, highlighting the need for normalization to subject-level means; (2) normalized qmALFF and qmfALFF are highly correlated with normalized amALFF and amfALFF, respectively, at both the subject and voxel levels; (3) *z*-scoring the BOLD signal does not affect amfALFF or qmfALFF, but it substantially alters amALFF and qmALFF. Based on these findings, we present a comprehensive flowchart of the (f)ALFF algorithm implemented in the temporal domain. The full procedure is implemented in R, and the corresponding script is available at: https://github.com/lejianhuang/ALFF.

## 1. Introduction

The amplitude of low-frequency fluctuations (ALFF) and its closely related measure, fractional amplitude of low-frequency fluctuations (fALFF), are two widely used resting-state fMRI analysis techniques for quantifying the “amount” of spontaneous neural activity within a specific frequency range [1–3]. These measures have been extensively applied to investigate brain activity in various clinical conditions [2, 4–6].

Despite their extensive use, inconsistencies in the definition and implementation of ALFF have led to the conflation of distinct mathematical constructs within the neuroimaging literature. First, the original description of ALFF is unclear, *i.e*., the average squared root of power instead of amplitude itself in the frequency domain [2]. Although the authors directly link the ALFF to amplitude in their updated software, the ambiguity has led to misrepresentation and errors in search results on platforms like Google and ChatGPT, which erroneously define ALFF as the total amplitude (sum) within a specified bandwidth. Second, variations in methodological implementations have also contributed to the conflation within the literature [7–10]. For instance, in one commonly used ALFF toolbox, RESTplus [7, 10], ALFF is computed as the arithmetic mean of the power spectrum amplitude within a given frequency range. In contrast, another popular software, Conn [8], indirectly defines ALFF using the quadratic mean of the amplitude. In another toolbox, BRANT [9], while its related article defines ALFF as the average square root of the power spectral density, its implementation is the same as REST [7] (the original version of RESTplus): the average square root of the power spectrum. Additionally, some researchers mistakenly computed ALFF after *z*-scoring the original BOLD signal, as is commonly done before calculating brain connectivity. Given these inconsistencies, it is imperative to rigorously revisit ALFF and fALFF.

In this paper, we mathematically clarify ALFF and fALFF in terms of the arithmetic mean (amALFF/amfALFF) and the quadratic mean (qmALFF/qmfALFF). We also reexamine the relationships between mean BOLD intensity, amALFF, amfALFF, qmALFF, and qmfALFF at both subject and voxel levels across two independent datasets on two different brain templates. Finally, we show how *z*-scoring the original BOLD signal affects ALFF and fALFF. Based on our findings, we present a flowchart illustrating the complete (f)ALFF algorithm implemented in the temporal domain.

## 2. Methods

### 2.1. Arithmetic Mean of ALFF

Let *s*(*m, n*) represent the preprocessed BOLD time-series signal of a voxel *m* at coordinates (*x, y, z*) in the brain after demeaning, where *m =* (*1, 2, …, M*) and *n =* (*0, 1, …, N − 1*). Here, *M* and *N* denote the number of voxels masked by a given brain template and volumes (time points) of the resting-state (RS) fMRI data, respectively. Let *s*_*bp*_(*m, n*) denote the BOLD signal after applying a bandpass temporal filter. In this study, we used a zero-lag fourth-order Butterworth filter with a frequency band of 0.01–0.08 Hz. For simplicity, voxel coordinates will be omitted in the following descriptions.

The amplitude spectrum of *s*(*m, n*) can be computed using the fast Fourier transform (FFT). The amplitude spectrum of *s*(*m, n*) before scaling is given by

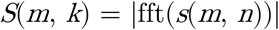

where *k =* (*0, 1, …, N − 1*) and the frequency interval between consecutive *k*-values is 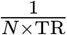 Hz. *TR* (repetition time) represents the time interval between consecutive volume acquisitions in the RS fMRI scan. For simplicity, and without loss of generality, we assume the number of sampling points, *N*, is even. The notation | · | represents the magnitude of each frequency component, computed as the square root of the sum of the squares of its real and imaginary parts after applying the FFT. *S*(*m, k*), *w*here *k =* (*1, 2, …, N − 1*), is symmetric around *k = N/*2.

Thus, the arithmetic mean of ALFF (amALFF) for a given voxel *m* within a specified frequency range [*f*_*l*_, *f*_*h*_] can be computed as

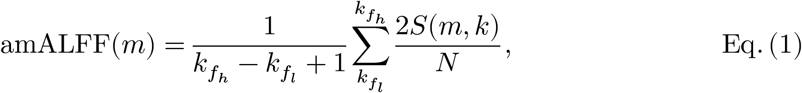

where *f*_*l*_ and *f*_*h*_ represent the lowest and highest frequencies in the specified range (set to 0.01 Hz and 0.08 Hz in this study), and 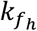 and 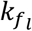 denote the FFT indices corresponding to or closest to *f*_*l*_ and *f*_*h*_, respectively. *N* is a scaling factor for normalization. This arithmetic mean of ALFF is consistent with the original definition and its derivative used for measuring ALFF in [2].

The arithmetic-mean fractional amplitude of low-frequency fluctuation (amfALFF) is defined as the ratio of the sum of amplitude spectrum within a specified frequency range [*f*_*l*_, *f*_*h*_] to that of the frequency range [*0, f*_*c*_]

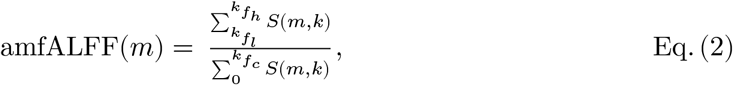

where *f*_*c*_ represents the low-pass cut-off frequency in the preprocessing stage (set to 0.2 Hz in this study), and 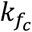 denotes the FFT index corresponding to or closest to *f*_*c*_.

### 2.2. Quadratic Mean of ALFF

The BOLD signal is an indirect measure of brain electrical activity. The mean power of *s*(*m, n*) represents the average energy of consumption per unit time for a voxel *m* and is calculated as

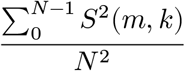

Accordingly, the mean power of *s*(*m, n*) within a specified frequency range [*f*_*l*,_ *f*_*h*_] is given by

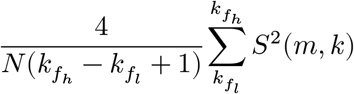

The corresponding mean amplitude, defined as the quadratic mean of ALFF (qmALFF)

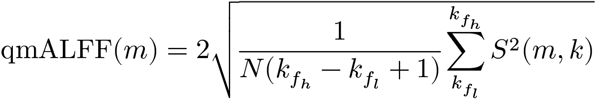

which can also be expressed as

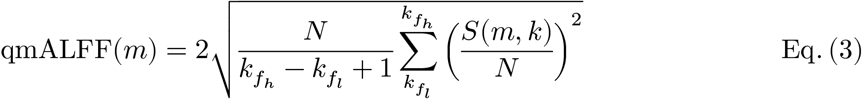

and similarly, the quadratic-mean fractional amplitude of low-frequency fluctuation (qmfALFF) can be given by

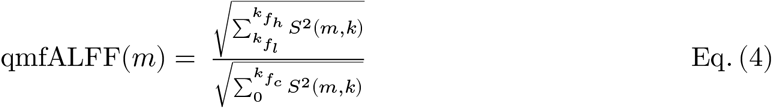

Based on Parseval’s Theorem, equation (3) can be rewritten in the form of time domain as

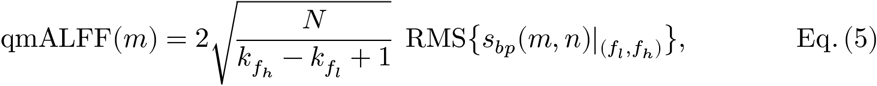

where 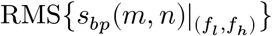 denotes the root mean square of *s*(*m, n*) filtered by a bandpass filter with cutoff frequencies *f*_*l*_ and *f*_*h*_. Thus, the qmfALFF in the form of time domain is given by

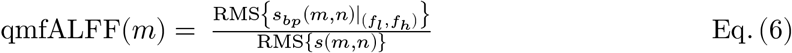

Note that *s*(*m, n*) has been filtered by a low-pass temporal filter with the low-pass cut-off frequency *f*_*c*_ in the preprocessing stage.

### 2.3. Correlations between mean BOLD intensity and am/qm(f)ALFF

From Equations (1) and (3), both amALFF and qmALFF across voxels are influenced by the number of volumes and sampling points (*N*); repetition time(*TR*), which determines FFT index (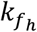 and 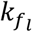) based on *f*_*l*_ and *f*_*h*_; and the mean BOLD intensity (MBI) across time series through *S*(*m, k*), even though *s*(*m, n*) is demeaned before calculating am/qmALFF. In a given study, *TR* and *N* remain constant once the study protocol is set. However, the MBI may vary due to several factors, such as scanner settings, participant physiology, and preprocessing steps [11, 12]. Therefore, it is essential to examine the correlations between MBI and am/qm(f)ALFF at both subject (mean of MBI across voxels) and voxel levels (MBI of voxel). Salient correlations that exist between these variables should be addressed before proceeding with further analyses. At subject level, the MBI is calculated as the mean of MBI across voxels within a given template in brain. At voxel level, the MBI is calculated as the mean of MBI across subjects for each voxel within a given template in a brain.

### 2.4. Normalization of am/qmALFF at the subject level

If a salient correlation exists between MBI and am/qm ALFF across subjects, it is reasonable to normalize the am/qm ALFF value to ensure the mean am/qm ALFF for each subject equals 1. This normalization minimizes inter-subject variability that is attributable to MBI and ensures comparability across individuals.

The normalized amALFF, denoted as amALFF_norm_, for the voxel *m* is calculated as

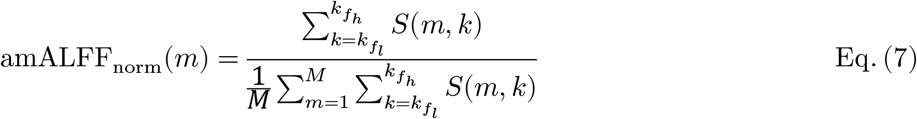

Equation (7) shows that the final analysis results in studies relying on the original version of ALFF (incorrectly defined as 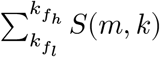) will not be affected if amALFF is normalized. The normalization allows the factors in equation (1) to be canceled out from the numerator and denominator of the equation (7).

Similarly, the normalized qmALFF, qmALFF_norm_, is defined as

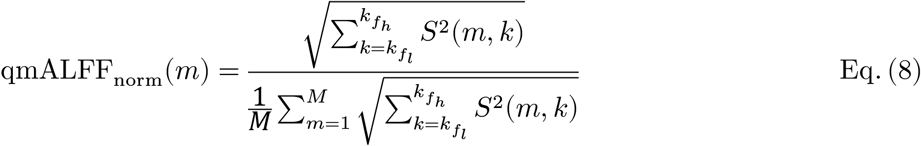

or

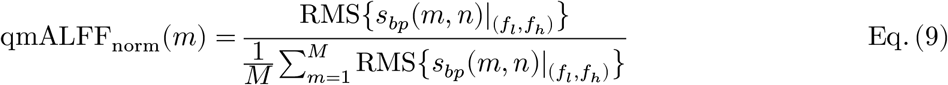

### 2.5. The effect of *z*-scoring on am/qm fALFF and am/qmALFF_norm_

*z*-scoring the BOLD signal over time does not alter the relative frequency content or the overall spectral shape. Instead, it scales the entire amplitude spectrum by a constant factor and removes the zero-frequency (demean). Mathematically, the *z*-scored amplitude spectrum is given by

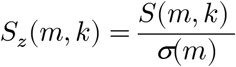

where *σ*(*m*) denotes the standard deviation of *s*(*m, n*), and *S*_*z*_(*m, k*) is the amplitude spectrum of the *z*-scored signal *s*(*m, n*). From equations (2) and (4), it follows that *z*-scoring does not affect amfALFF or qmfALFF, since *σ*(*m*) in each equation can be canceled out from the numerator and denominator. Moreover, according to equation (6), when *s*(*m, n*) is *z*-scored, such that *RMS{s*(*m, n*)*} = 1*, qmALFF becomes

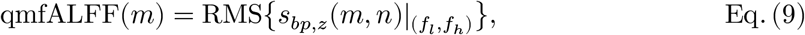

where 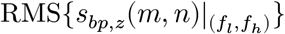 represents the root mean square of *z*-scored signal *s*(*m, n*), filtered using a bandpass filter with cutoff frequencies *f*_*l*_ and *f*_*h*_.

By *z*-scoring the normalized metrics, amALFF_norm_ and qmALFF_norm_, we obtain

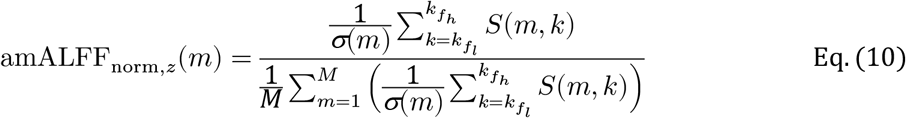

and

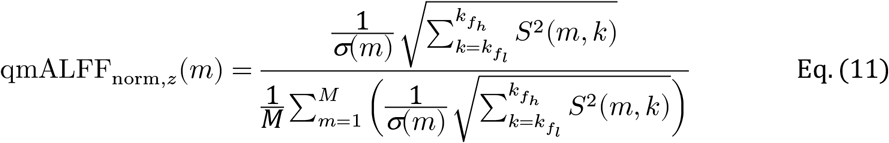

These expressions indicate that amALFF_norm_ and qmALFF_norm_ are equal to their *z*-scored counterparts (amALFF_norm,*z*_ and qmALFF_norm,*z*_) if and only if *σ*(1) *= σ*(2) *=* … *= σ*(*M*).

### 2.6 Datasets

This study utilizes two datasets. The first dataset, from a previous study in China, includes 157 healthy controls in [13] (China157) (77 males, 80 females; mean age ± SD: 40 ± 14 years old).

The second dataset comes from the Human Connectome Project 1200 (HCP1200) [14], which includes 926 young healthy volunteers after quality control (425 males, 501 females; mean age m ± SD: 19 ± 4 years old). We found that the HCP1200 dataset has three clusters of MBI (subjects with low (*l*), medium (*m*), and high (*h*) MBI are 14, 759, and 153, respectively).

### 2.7 MRI scanning parameters

For the China157 dataset, subjects were scanned on a 3 Tesla GE-Discovery 750. T1 images were acquired with following parameters: voxel size = 1 × 1 × 1 mm^3^; TR/TE = 7.7/3.4 ms; flip angle = 12°; field of view = 256 × 256 mm^2^. RS-fMRI images were acquired with the following parameters: voxel size = 3.4375 × 3.4375 × 3.5 mm^3^; TR/TE = 2500/30 ms; flip angle = 90°; in-plane resolution = 64 × 64; field of view = 220 × 220 mm^2^; number of volumes = 230; slices per volume = 42, which covers the whole brain from the cerebellum to the vertex.

For the HCP1200 dataset, subjects were scanned on a 3 Tesla GE-Discovery 750. T1 images were acquired with following parameters: voxel size = 0.7 × 0.7 × 0.7 mm^3^; TR/TE = 2400/2.14 ms; flip angle = 8°; field of view = 224 × 224 mm^2^. RS-fMRI images were acquired with the following parameters: voxel size = 2 × 2 × 2 mm^3^; TR/TE = 720/33.1 ms; flip angle = 52°; field of view = 208 × 180 mm^2^; number of volumes = 1200; multiband accelerator = 8; slices per volume = 72, which covers the whole brain from the cerebellum to the vertex.

### 2.8. RS-fMRI data preprocessing and denoising

Both datasets were preprocessed using fmriprep version of 23.1.4 [15], a Nipype based tool [16, 17]. Each T1w (T1-weighted) volume was skull-stripped using antsBrainExtraction.sh v2.1.0 (using the OASIS NKI template). Brain surfaces were reconstructed using recon-all from FreeSurfer v6.0.1 [18], and the brain mask estimated previously was refined with a custom variation of the method to reconcile ANTs-derived and FreeSurfer-derived segmentations of the cortical gray-matter of Mindboggle [19]. Spatial normalization to the ICBM 152 Nonlinear Asymmetrical template version 2009c [20] was performed through nonlinear registration with the antsRegistration tool of ANTs v2.1.0 [21], using brain-extracted versions of both T1w volume and template. Brain tissue segmentation of cerebrospinal fluid (CSF), white-matter (WM), and gray-matter (GM) was performed on the brain-extracted T1w using fast [22].

Functional data was slice time corrected using 3dTshift from AFNI v16.2.07 [23] and motion corrected using mcflirt [24]. No distortion correction was performed. This was followed by co-registration to the corresponding T1w using boundary-based registration [25] with six nine twelve degrees of freedom, using bbregister (FreeSurfer v6.0.1). flirt (FSL). Motion correcting transformations and BOLD-to-T1w transformation and T1w-to-template (MNI) warp were concatenated and applied in a single step using antsApplyTransforms (ANTs v2.1.0) using Lanczos interpolation.

Physiological noise regressors were extracted applying CompCor [26]. Principal components were estimated for the two CompCor variants: temporal (tCompCor) and anatomical (aCompCor). A mask to exclude signal with cortical origin was obtained by eroding the brain mask, ensuring it only contained subcortical structures. Six tCompCor components were then calculated including only the top 5% variable voxels within that subcortical mask. For aCompCor, six components were calculated within the intersection of the subcortical mask and the union of CSF and WM masks calculated in T1w space, after their projection to the native space of each functional run. Frame-wise displacement [27] was calculated for each functional run using the implementation of Nipype. ICA-based Automatic Removal Of Motion Artifacts (AROMA) was used to generate aggressive noise regressors as well as to create a variant of the data that is non-aggressively denoised [28].

During the denoising stage, a confound regressor matrix was used to regress out confounding effects from the preprocessed RS-fMRI data, which included six head-motion parameters, the average signal within anatomically derived eroded WM mask, the average signal within anatomically derived eroded CSF mask, the average signal within the brain mask (Global), non-steady-state-outliers (to remove outliers in the time series), and discrete cosine-basis regressors (to correct low-frequency signal drift). Finally, a 4th-order Butterworth low-pass temporal filter was applied, with a cutoff frequency of 0.2 Hz.

## 3. Results

### 3.1. Subjects with greater mean BOLD intensity have greater amALFF and qmALFF but not amfALFF or qmfALFF

At the subject level, MBI is calculated as the mean of MBI across all voxels within a given template in brain. As shown in **Fig. 1a** (China157) and **Fig. 1b** (HCP1200), at the subject level, MBI positively correlates amALFF in both the China157 (*p* = 0.02, *r* = 0.18) and HCP1200 (*p* < 0.001, *r* = 0.91) datasets. Similarly, MBI correlates with qmALFF (China157: *p* = 0.02, *r* = 0.18; HCP1200: *p* < 0.001, *r* = 0.91). These results underscore the necessity of normalizing amALFF and qmALFF by dividing them by their mean across all voxels within a specified mask to minimize inter-subject variability attributable to MBI and ensures comparability across subjects. The normalized amALFF and qmALFF are denoted as amALFF_norm_ and qmALFF_norm_, respectively, ensuring that their mean equals 1.

**Figure 1.**
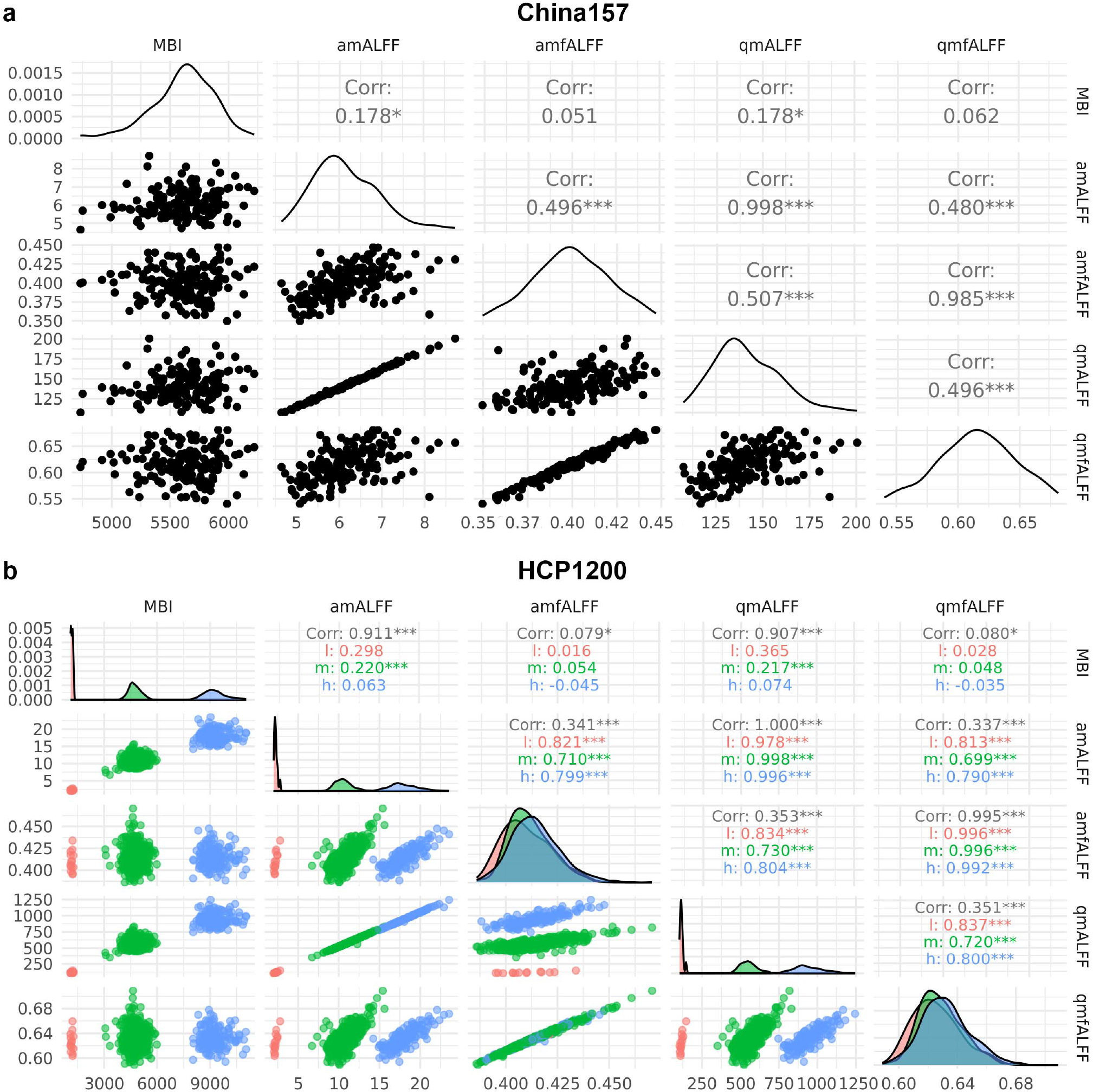
Correlation matrix of mean BOLD intensity (MBI), amALFF, amfALFF, qmALFF, and qmfALFF for the China157 and HCP1200 datasets at the subject level. **a)** For the China157 dataset, the MBI is weakly but significantly correlated with amALFF (*p* = 0.02, *r* = 0.178) and qmALFF (*p* = 0.02, *r* = 0.178) but not significantly correlated with amfALFF (*p* = 0.51, *r* = 0.051) or qmfALFF (*p* = 0.43, *r* = 0.062), Additionally, amALFF is significantly correlated with qmALFF (*p* < 0.001, *r* = 0.998), and amfALFF is significantly correlated with qmfALFF (*p* < 0.001, *r* = 0.995). The correlations between amALFF and amfALFF (*p* < 0.001, *r* = 0.496) and between qmALFF and qmfALFF (*p* < 0.001, *r* = 0.496) are significant. **b)** Red, green, and blue colors represent 3 clusters with low (l), medium (m), and high (h) MBI value, respectively, for the HCP1200 dataset. As a whole, the MBI is significantly correlated with amALFF (*p* < 0.001, *r* = 0.911), qmALFF (*p* < 0.001, *r* = 0.907), amfALFF (*p* = 0.02, *r* = 0.079), and qmfALFF (*p* = 0.02, *r* = 0.080), Additionally, amALFF is significantly correlated with qmALFF (*p* < 0.001, *r* = 1), and amfALFF is significantly correlated with qmfALFF (*p* < 0.001, *r* = 0.995). The correlations between amALFF and amfALFF (*p* < 0.001, *r* = 0.341) and between qmALFF and qmfALFF (*p* < 0.001, *r* = 0.351) are significant. However, in the individual cluster, only the medium MBI cluster is significantly correlated with amALFF (*p* < 0.001, *r* = 0.22), and qmALFF (*p* < 0.001, *r* = 0.22). The correlations are greatly increased both between amALFF and amfALFF (l: 0.821, m: 0.710, and h:0.799) and between qmALFF and qmfALFF (l: 0.813, m: 0.699, and h:0.79).

In contrast, MBI did not consistently scale with amfALFF (China157: *p* = 0.51, *r* = 0.05; HCP1200: low (l, pink): *p* =0.96, *r* = 0.02; medium (m, green): *p* = 0.14, *r* = 0.05; high (h, blue): *p* = 0.58, *r* = 0.02) or qmfALFF (China157: *p* = 0.43, *r* = 0.06; HCP1200: low (l, pink): *p* =0.92, *r* = 0.03; medium (m, green): *p* = 0.18, *r* = 0.05; high (h, blue): *p* = 0.67, *r* = −0.04)).

### 3.2. Across-subject Relationships between amALFF, qmALFF, amfALFF, and qmfALFF

As shown in **Fig. 1a** (China157) and **Fig. 1b** (HCP1200), qmALFF and amALFF appear to be linear transformations of one another in both the China157 (*p* < 0.001, *r* = 1.00) and HCP1200 (*p* < 0.001, *r* = 1.00 across all clusters) datasets. Similarly, so too are qmfALFF and amfALFF (China157: *p* < 0.001, *r* = 1.00; HCP1200: *p* < 0.001, *r* = 1.00 across all clusters). When performing full-brain ALFF analyses across individuals, qm(f)ALFF and am(f)ALFF yield similar results.

Additionally, amALFF and amfALFF are positive correlated in both the China157 (*p* < 0.001, *r* = 0.50) and HCP1200 datasets (low (l, pink): *p* < 0.001, *r* = 0.82; medium (m, green): *p* < 0.001, *r* = 0.71; high (h, blue): *p* < 0.001, *r* = 0.80)). A similar pattern is observed the correlations between qmALFF and qmfALFF (China157: *p* < 0.001, *r* = 0.51 and HCP1200: low (l, pink): *p* < 0.001, *r* = 0.83; medium (m, green): *p* < 0.001, *r* = 0.73; high (h, blue): *p* < 0.001, *r* = 0.80).

### 3.3. Mean BOLD intensity is associated with amfALFF and qmfALFF but not with amALFF_norm_ or qmALFF_norm_ at the voxel level

At voxel level, the MBI is calculated as the mean of MBI across subjects for each voxel within a given template in a brain. As shown in **Fig. 2d** (China157 with MNI152 brain mask) and **Fig. 2f** (China157 with Schaefer mask), and **Fig. 2e** (HCP1200 with MNI152 brain mask) and **Fig. 2g** (HCP1200 with Schaefer mask), at the voxel level, MBI is moderately correlated with amfALFF in both the China157 (*p* < 0.001, *r* = 0.23; *p* < 0.001 *r* = 0.25) and HCP1200 (*p* < 0.001, *r* = 0.21; *p* < 0.001 *r* = 0.26) datasets for both masks. Similarly, MBI is significantly moderately correlated with qmfALFF (China157: *p* < 0.001, *r* = 0.23 and *p* < 0.001, *r* = 0.26; HCP1200: *p* < 0.001, *r* = 0.22 and *p* < 0.001, *r* = 0.26). However, MBI’s correlation with the normalized metrics, amALFF_norm_ or qmALFF_norm_, is close to 0. At both the subject and the voxel levels, MBI was nearly orthogonal to the normalized ALFF metrics.

**Figure 2.**
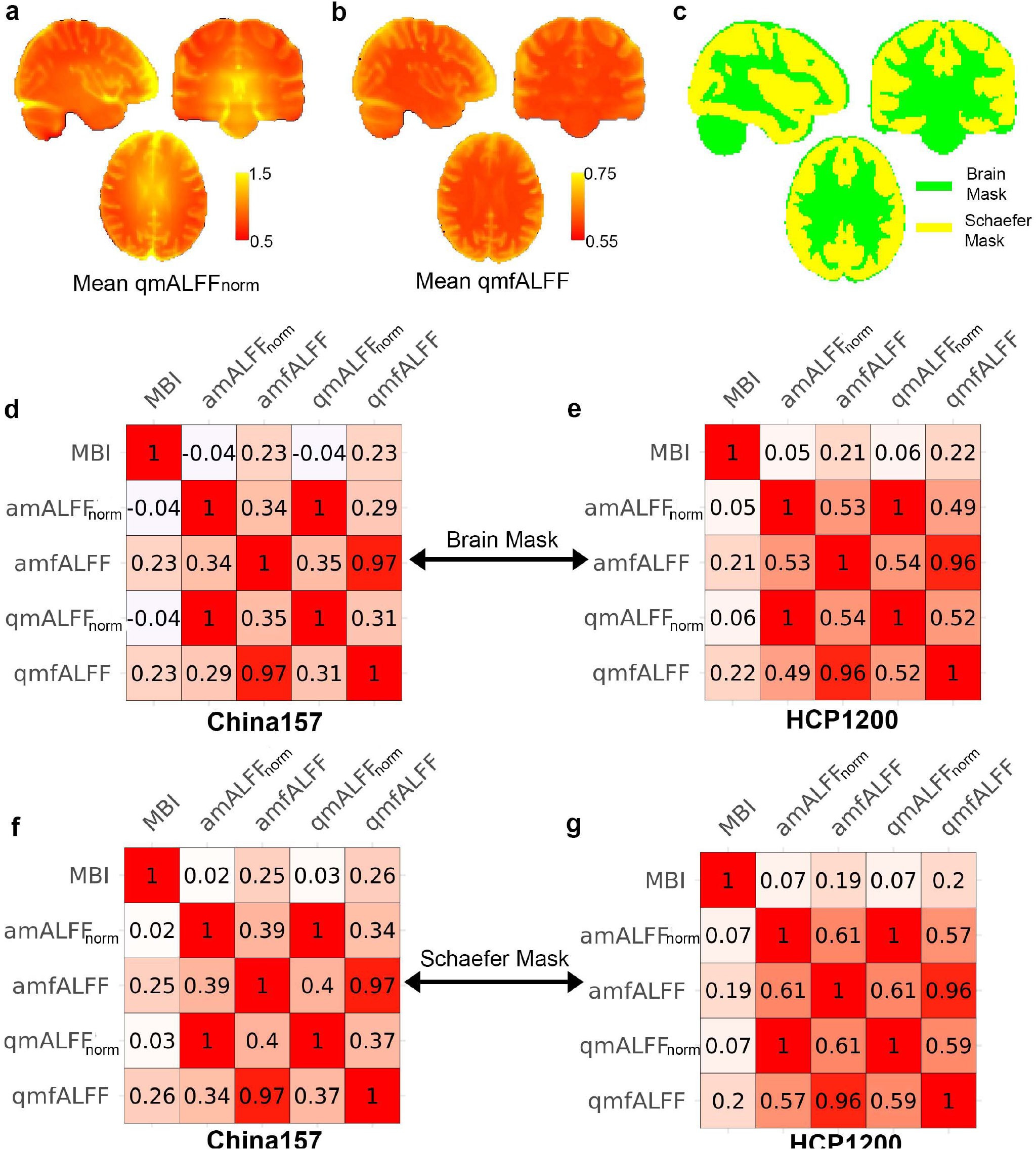
Correlation matrix of mean BOLD intensity (MBI), amALFF_norm_, amfALFF, qmALFF_norm_, and qmfALFF for the China157 and HCP1200 datasets at the voxel level. **a)** and **b)** Visualization of the mean qmALFF_**norm**_ and qmfALFF_**norm**_ across subject in the HCP1200 dataset. **c)** Illustration of the MNI152 brain (green) and Schaefer (yellow) mask used in the analysis. **d)** In the China157 dataset on the MNI152 brain mask, the MBI is significantly correlated with amfALFF (*p* < 0.001, *r* = 0.23) and qmfALFF (*p* < 0.001, *r* = 0.23), but shows no significant correlation with amALFF_**norm**_ or qmALFF_**norm**_. Notably, amALFF_**norm**_ is highly significantly correlated with qmALFF_**norm**_ (*p* < 0.001, *r* = 1), and amfALFF is significantly correlated with qmfALFF (*p* < 0.001, *r* = 0.97). Additionally, the relationships between amALFF_**norm**_ and amfALFF (*p* < 0.001, *r* = 0.34), as well as between qmALFF_**norm**_ and qmfALFF (*p* < 0.001, *r* = 0.31), show moderate but statistically significant correlations. **e)** In the HCP1200 dataset on the MNI152 brain mask, the MBI is significantly correlated with amfALFF (*p* < 0.001, *r* = 0.21) and qmfALFF (*p* < 0.001, *r* = 0.22), but shows no significant correlation with amALFFN or qmALFF_**norm**_. Notably, amALFF_**norm**_ is highly significantly correlated with qmALFF_**norm**_ (*p* < 0.001, *r* = 1), and amfALFF is significantly correlated with qmfALFF (*p* < 0.001, *r* = 0.96). Additionally, the relationships between amALFF_**norm**_ and amfALFF (*p* < 0.001, *r* = 0.53), as well as between qmALFF_**norm**_ and qmfALFF (*p* < 0.001, *r* = 0.52), show moderate but statistically significant correlations. **f)** In the China157 dataset on the Schaefer mask, the MBI is significantly correlated with amfALFF (*p* < 0.001, *r* = 0.25) and qmfALFF (*p* < 0.001, *r* = 0.26), but shows no significant correlation with amALFF_**norm**_ or qmALFFN. Notably, amALFFN is highly significantly correlated with qmALFFM (*p* < 0.001, *r* = 1), and amfALFF is significantly correlated with qmfALFF (*p* < 0.001, *r* = 0.97). Additionally, the relationships between amALFF_**norm**_ and amfALFF (*p* < 0.001, *r* = 0.39), as well as between qmALFF_**norm**_ and qmfALFF (*p* < 0.001, *r* = 0.37), show moderate but statistically significant correlations. **g)** In the HCP1200 dataset on the Schaefer mask, the MBI is significantly correlated with amfALFF (*p* < 0.001, *r* = 0.19) and qmfALFF (*p* < 0.001, *r* = 0.20), but shows no significant correlation with amALFF_**norm**_ or qmALFF_**norm**_. Notably, amALFF_**norm**_ is highly significantly correlated with qmALFFN (*p* < 0.001, *r* = 1), and amfALFF is significantly correlated with qmfALFF (*p* < 0.001, *r* = 0.96). Additionally, the relationships between amALFFN and amfALFF (*p* < 0.001, *r* = 0.61), as well as between qmALFFN and qmfALFF (*p* < 0.001, *r* = 0.59), show moderate but statistically significant correlations.

### 3.4. Relationships among amALFF_norm_, qmALFF_norm_, amfALFF, and qmfALFF at the voxel level

As shown in **Fig. 2d** (China157 with MNI152 brain mask), **Fig. 2f** (China157 with Schaefer mask), **Fig. 2e** (HCP1200 with MNI152 brain mask), and **Fig. 2g** (HCP1200 with Schaefer mask), across voxels, qmALFF_norm_ strongly correlated with amALFF_norm_ across both datasets and masks (China157: *p* < 0.001, *r* = 1.00 for both masks; HCP1200: *p* < 0.001, *r* = 1.00 for MNI152 brain mask and *p* < 0.001, *r* = 0.96 for Schaefer mask). Similarly, qmfALFF strongly correlated with amfALFF (China157: *p* < 0.001, *r* = 0.97 for both masks; HCP1200: *p* < 0.001, *r* = 0.97 for both masks), indicating a strong relationship between the two variants of ALFF across voxels, as also observed across subjects.

Furthermore, across voxels, amALFF_norm_ and amfALFF exhibit modest correlations in both China157 (*p* < 0.001, *r* = 0.34 for brain mask and *p* < 0.001, *r* = 0.39 for Schaefer mask) and HCP1200 datasets (*p* < 0.001, *r* = 0.53 for brain mask and *p* < 0.001, *r* = 0.61 for Schaefer mask). A similar pattern is observed in the correlations between qmALFF_norm_ and qmfALFF (China157: *p* < 0.001, *r* = 0.29 for brain mask and *p* < 0.001, *r* = 0.34 for Schaefer mask, and HCP1200: *p* < 0.001, *r* = 0.49 for brain mask and *p* < 0.001, *r* = 0.57 for Schaefer mask). As shown in **Fig. 2a** and 2**b**, correlations between am(qm)ALFF_norm_ and am(qm)fALFF are consistently higher when using the Schaefer mask compared to the brain mask. This is likely due to the presence of elevated am(qm)ALFF_norm_ values in white matter and vascular regions, which contribute to greater dissimilarity between the normalized and fractional metrics on the whole-brain mask.

### 3.5. *z*-scoring disrupts the distribution of qmALFF_norm_

From equations (9) and (10), it is apparent that *z*-scoring the BOLD signal disrupts both qmALFF_norm_ and amALFF_norm_ across voxels. The extent of this disruption depends on the standard deviation of the BOLD signal at each voxel. **Fig.3a** and **3b** illustrate the nonlinear relationship between qmALFF_norm_ and qmALFF_*z*,norm_, along with corresponding histograms of both metrics for the averaged China157 and HCP1200 datasets with Schaefer mask, respectively. In the China157 dataset, qmALFF_*z*,norm_ shares only 4% of the variance with qmALFF_norm_, while in the HCP1200 dataset, the shared variance is 28%. These findings indicate that z-scoring significantly alters the distribution of qmALFF_norm_; thus, using *z*-scored BOLD signal to compute ALFF is inappropriate.

**Figure 3.**
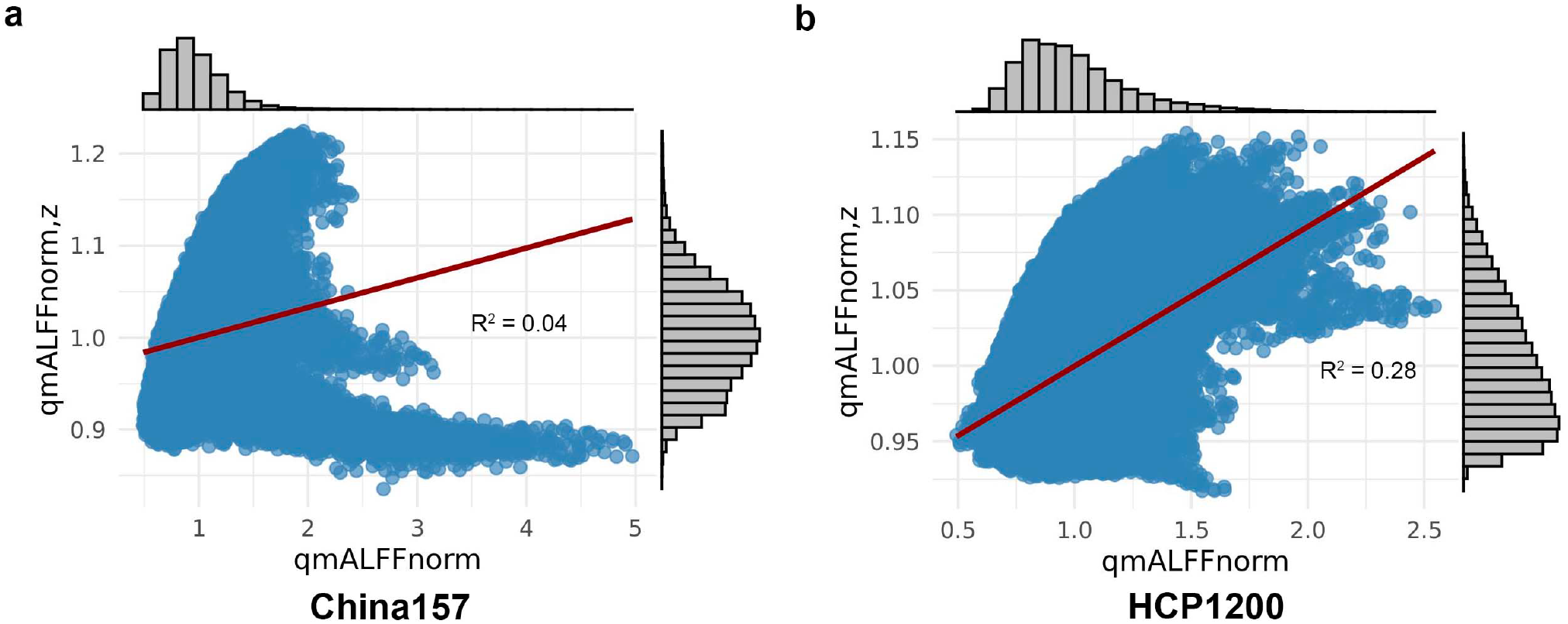
Scatter plots illustrating the nonlinear relationship between qmALFF_norm_ and qmALFFZ_norm,z_, along with corresponding histograms of both metrics. **a)** *R*^2^ = 0.04 for China157 dataset with Schaefer mask; **b)** *R*^2^ = 0.28 for HCP1200 dataset with Schaefer mask.

### 3.4. Procedure for calculating qmALFF_norm_ and qmfALFF

**Fig. 4** illustrates the complete procedure for calculating qmALFF_norm_ and qmfALFF, beginning with the preprocessing stage. The preprocessed RS-fMRI data is subjected to temporal band-pass filtering using a zero-lag fourth-order Butterworth filter with the frequency band of 0.01–0.08 Hz. Subsequently, qmALFF is calculated according to equation (5) and qmALFF_norm_ by normalizing qmALFF with its mean value. Simultaneously, qmfALFF is computed using equation (6). All calculations are performed in the time domain, without applying an FFT. This procedure is implemented in R, and the corresponding script is available on https://github.com/lejianhuang/ALFF.

**Figure 4.**
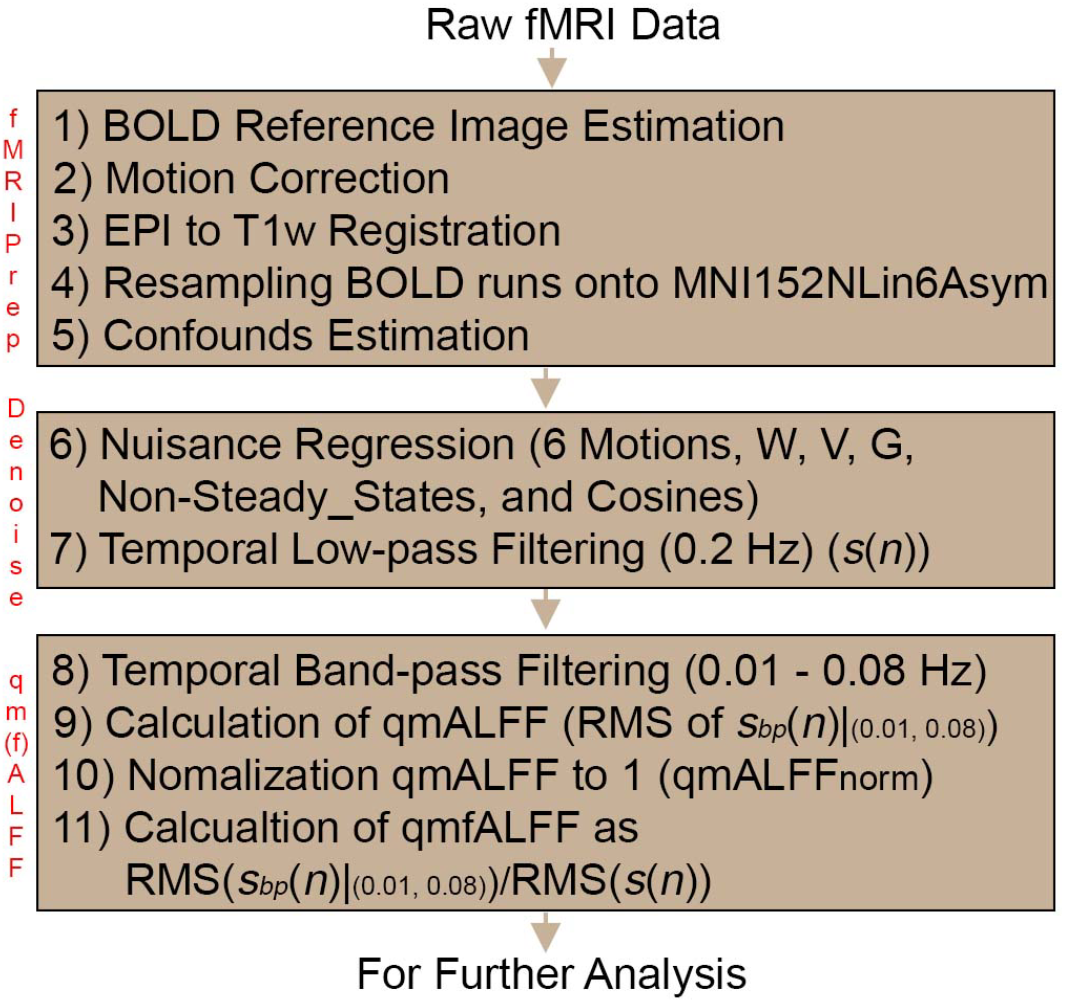
Flowchart of calculating qmALFF_norm_ and qmfALFF, starting from the preprocessing stage.

## Discussion

In this study, we clarified and resolved the inconsistencies in the definition and implementation of ALFF, and we introduced a novel variant called qmALFF. Our key findings include: (1) The mean BOLD intensity is strongly associated with both amALFF and qmALFF, necessitating normalization to their respective mean MBI across voxels; (2) qmALFF_norm_ and qmfALFF are highly correlated with amALFF_norm_ and amfALFF, respectively, at both subject and voxel levels; (3) *z*-scoring the BOLD signal does not affect amfALFF or qmfALFF, but it substantially alters both amALFF and qmALFF.

The confusion surrounding the calculation of ALFF originates from the original publication by [2]. The authors stated: “*Since the power of a given frequency is proportional to the square of the amplitude of this frequency component […], the square root was calculated at each frequency of the power spectrum and the averaged square root was obtained […]. This averaged square root was taken as the ALFF*” (Section of 2.5. ALFF analysis in by [2]). They used the SPM software [29] to compute the square of the amplitude at each frequency component, followed by averaging their square roots – an indirect and potentially misleading process for calculating the arithmetic mean of ALFF. This approach introduces ambiguity in the field, blurring the conceptual line between “amplitude” and “mean amplitude”. More precisely, the mean amplitude of low-frequency fluctuation reflects changes in spontaneous neuronal activity. Although the normalizing amALFF by its subject-level mean (i.e., computing amALFF_norm_) effectively removes the effect of MBI and reduces inter-subject variability across individuals, it also masks the definitional inconsistency that persists in sources such as Google or ChatGPT –namely, defining ALFF as the sum of amplitudes rather than the mean amplitude. This discrepancy becomes mathematically obscured because the factors in Equation (7) cancel out during normalization. Fortunately, our results demonstrate that these definitional inconsistencies do not compromise previous findings. Our data clearly show that amALFF_norm_ and qmALFF_norm_ are nearly perfectly correlated (r ≈ 1) at both subject and voxel levels across two datasets and two different mask templates.

Our proposed qmALFF method, which derives from the concept of signal energy and its corresponding average power, provides a more physically grounded measure of mean amplitude in the frequency domain. In physics and engineering, the power or energy of a signal is proportional to the square of its amplitude – a principle to which qmALFF directly relates. Importantly, qmALFF can be computed either in the frequency domain (equation (3)) or in the time domain (equation (5)), making it flexible for implementation. Notably, the Conn toolbox (https://web.conn-toolbox.org/fmri-methods/connectivity-measures/other) [8] computes ALFF and fALFF using the time-domain formulation, but incorrectly cites references [2, 3], which describe the original and potentially flawed definitions of ALFF and fALFF. Our work helps reconcile this inconsistency.

qmfALFF, like amfALFF, quantifies the relative contribution of low-frequency fluctuations within a given band to the whole frequency range determined by the cut-off frequencies in the preprocessing stage [3, 30]. Although the correlation between qmfALFF and qmALFF is moderate (r = 0.3–0.5, see **Fig. 1**), qmfALFF has a key advantage: it is insensitive to MBI at the subject level and unaffected by *z*-scoring of the BOLD. As such, it remains an essential measure for assessing the intensity of spontaneous neural activity within a specified frequency band. However, at the voxel level, qmfALFF shows a modest association with MBI (r ≈ 0.2, see **Fig. 2**). Addressing how to remove this voxel-level MBI effect in qmfALFF remains an open question for future work.

Several factors influence MBI. First, individual differences—such as variations in brain structure, vascularization, and metabolism—can lead to variability across subjects [11, 12]. Second, physiological noise during scanning, including respiration, cardiac activity, and brain motion, which is strongly linked to respiration [31], can affect MBI measurements. Third, scanner settings play a significant role. Factors such as manufacturer, magnetic field strength, echo time, repetition time, and flip angle all impact the MBI. In this study, the two datasets analyzed were acquired using scanners from different manufacturers, with identical magnetic field strengths but differing echo times, repetition times, and flip angles. Additionally, the scanner software version is another important variable. For instance, subjects in the HCP1200 dataset were scanned using the same brand of scanner with consistent parameters over three years [14, 32]. However, as shown in **Fig. 1b**, three distinct clusters of MBI values are observed. A plausible explanation is that the scanner software was updated three times during this period, potentially altering data resolution and applying GE’s auto-scaling processes, which constrain the data range. Although the signal-to-noise ratio remained relatively stable, the MBI varied significantly, which in turn affected the am/qmALFF values. Therefore, normalizing am/qmALFF is essential to reduce inter-subject variability caused by these factors and to ensure comparability across individuals.

## Limitations

The use of multiple sites or centers in clinical trials and data-sharing initiatives to expand the subject pool has become a common practice. However, we observed that the distribution of am(qm)fALFF values, as shown in **Fig. 1**, differs significantly between the two datasets analyzed. This discrepancy highlights the need to address site- and dataset-specific effects. To enable unbiased comparisons of ALFF or fALFF across different populations, it is essential to develop a robust framework that accounts for these inter-site and inter-dataset variations.

## Conclusions

In this study, we provided a mathematical clarification of ALFF and fALFF by introducing two novel variants: amALFF and qmALFF. Upon identifying an appreciable correlation between am(qm)ALFF and MBI, we emphasized the importance of normalizing these measures to subject-level means. Compared to amALFF, qmALFF offers the advantage of being computable directly in the time domain. To facilitate reproducibility and further research, we have implemented the complete procedure for calculating qm(f)ALFF in R. The corresponding script is publicly available at: https://github.com/lejianhuang/ALFF.

## Data available

The script is available in https://github.com/lejianhuang/ALFF. The HCP1200 dataset is downloaded from the Human Connectome Project: https://www.humanconnectome.org/study/hcp-young-adult/. The China157 dataset will be available in https://openpain.org once upon this manuscript is accepted for publication.

## Acknowledgement

We thank NIH (grant 1P50DA044121-01A1) for funding data analysis. This material is based upon work supported by the National Science Foundation Graduate Research Fellowship under Grant No. DGE-1324585.

